# Survey of transcription initiation in the streamlined genomes of *Paramecium*

**DOI:** 10.64898/2026.09.19.752804

**Authors:** Berenice Jiménez-Marín, Darian Stickling, Tia Swenty, Jean-François Goût, Samuel Miller, Michael Lynch

## Abstract

In the genus *Paramecium*, the macronuclear genome is remarkably compact and optimized for gene expression. As a means to explore eukaryotic transcription in the context of a streamlined genome and shed light on the role of sequence architecture on gene expression and loss, we analyzed the distribution and diversity of candidate transcription initiation sites (TISs) in *Paramecium sexaurelia, Paramecium tetraurelia* and their outgroup, *Paramecium caudatum.* Our analysis suggests that for *Paramecium,* most genes have very short 5’ UTRs (40 bp or less) and their transcription initiation regions (TIRs) have a median dispersion (akin to width) of 8-10 bp. The TIRs for the three species have high AT content. TIR dispersion is not to gene expression. However, mean TIS position relative to the translation start site per gene does in gene expression for the three species, and is often conserved between them. While mean TIS position and gene expression are linked, gene expression itself is the main driver of paralog retention in the *aurelias*. As compared to other eukaryotes, *Paramecium* has a uniquely well-defined and short main TIS region, and sequence motifs that likely diverge from the consensus in multicellular eukaryotes.

## Introduction

Transcriptional regulation is a key process underlying the maintenance of organismal function. Intrinsic cues, such as developmental and homeostatic signals, and extrinsic cues, such as environmental challenges, result in an orchestrated response through precise regulation of distinct genes. Regulation of gene expression stems from the intricate coordination of players, among them the core promoter. The core promoter is usually a short DNA region upon which transcription factors (TFs) and RNA polymerase II are recruited to assemble the preinitiation complex (Roeder, 1996, Roy and Singer, 2015), such that transcription initiates downstream of it, often resulting in a 5’ untranslated region (5’ UTR). Certain core promoter sequences are evolutionarily conserved from bacteria to metazoans, such as the CCAAT box (Roy and Singer, 2015), whereas the TATA box and the initiator (Inr) sequence are conserved across eukaryotes (Lo and Smale, 1996, Schumacher et al., 2003, Yang et al., 2007, Villao-Uzho et al., 2007, Roy and Singer, 2015).

In bacterial genomes, the main RNA polymerase initiation factor variant, sigma 70, relies on a 6-bp sequence at position -35 (35 bases upstream of the transcription initiation site) that binds the RNA polymerase sigma factor to the DNA, while a 6-bp sequence at -10 facilitates the opening of the DNA to form the transcription bubble (Pribnow, 1975, Feklistov and Darst, 2011). The sequences at these positions and the length of the sequence between them impact the intensity of gene expression (Aoyama et al., 1983, Singh et al. 2011), as do additional sequences that modulate promoter strength (Chen et al. 2021). Other sigma factors recognize different sites, for example, sigma 54 with sites located at -24 and -12 (Danson et al. 2019). Furthermore, leaderless genes (with no 5’ UTR) have been found in *M. tuberculosis* (Cortes et al. 2014). Upon transcription initiation, the bacterial 5’ UTR can itself be involved in regulatory activity (Liu et al. 2024).

The eukaryotic core promoter is believed to be fairly information-rich and may include several TF recognition motifs (Juven-Gershon and Kadonaga 2010). In addition, core promoters serve as integrating centers for enhancer signals and have at least one binding site for an RNA polymerase. The interaction between these elements influences the transcriptional profile of their targets (Juven-Gherson et al. 2008, Zabidi and Stark, 2016). For protein-coding genes, a binding site for RNA polymerase II is necessary, but not sufficient, for promoter recognition. General transcription-factor binding sites are the key for transcription initiation site (TIS) recognition, and no one master TF is common to all promoters, at least in mammals and *Drosophila* (Sandelin et al. 2007, Kadonaga, 2012).

Surveys of transcription initiation in mice and humans suggest that initiation is not necessarily linked to a unique position in the core promoter but rather forms a region (TIR) that can have one or more preferred initiation sites (peaks) and may be focused (confined to a short stretch of contiguous nucleotides) or dispersed (Carninci et al. 2006). The focused/dispersed dichotomy for TIRs is present in *Drosophila* species (Main et al., 2013), as well as *Caenorhabditis elegans* (Chen et al. 2013), albeit in different proportions. In mammals, distinct TIR shapes appear to correspond to different promoter and functional-product classes (Carninci et al. 2006). Conversely, in non-vertebrates, transcription initiation is expected to be mostly linked to a focused single site of initiation (Juven-Gershon and Kadonaga 2010).

While *Saccharomyces cerevisiae* does have both focused and dispersed TIRs (Miura et al. 2006), some TIRs show greater dispersion as a result of the evolution of a scanning mechanism of transcription initiation, whereas other yeasts (such as *Schizosaccharomyces pombe*) display standard focused TIRs for TATA-box containing promoters (Lu and Lin, 2021). The TIRs of *Plasmodium falciparum* are overall dispersed, located within 500 bp upstream of the transcription start site for coding genes; however, most TIRs occur 50 bp upstream of the start codon (Adjalley et al., 2016). In *Trypanosoma brucei,* which has no recognizable promoters, TIRs are extremely dispersed, spanning a region of up to 2 kb (Wedel et al., 2017). The modular nature and combinatorial potential of core promoters have thus made the study of the evolution of gene expression circuits a challenging field and encourage further exploration with the simplest eukaryotic models possible.

The *Paramecium* ciliate model system consists of a group of species suitable for the study of promoter evolution, genetics, and speciation (Long et al. 2023). The genus contains approximately 50 species – all unicellular, heterotrophic, aquatic eukaryotes. Each *Paramecium* cell contains two nuclei, a transcriptionally inactive germline micronucleus, and an active, somatic macronucleus. The macronuclear genome is polyploid, AT rich, and heavily edited from its micronuclear source, such that it is streamlined for constant transcriptional activity (Amar 1994, McGrath et al. 2014, Long et al. 2023, Ni et al. 2025). *Paramecium* genes are, on average, 1.4 kb in length, and contain on average just two short (∼20-40 bp) introns (McGrath et al. 2014, Ni et al. 2025). Their intergenic regions are also compact, mostly between 40 and 700 bp long (McGrath et al. 2014, Arnaiz et al. 2017, Long et al. 2023, Ni et al. 2025).

Within the genus *Paramecium*, the *aurelia* cryptic species complex is thought to have arisen following at least two whole-genome duplications (WGD) (McGrath et al. 2014, Long et al. 2023, Ni et al. 2025). Gene-duplicate retention is very high in this lineage; *P. tetraurelia* has retained about 49.6% of its duplicated genes, and *P. sexaurelia* 41.6% (McGrath et al. 2014). The retention of distinct gene members per paralog set is correlated to the GC content per gene, the intensity of expression of each member gene, and the functional role of the paralogs (McGrath et al. 2014, Johri et al. 2017, Gout et al. 2023). Analyses performed on multiple *aurelia* species, as well as two outgroups (*P. caudatum and P. multimicronucleatum*) suggest that dosage is a key determinant of gene fate (Gout et al. 2023). It was proposed that the slow duplicate loss in *Paramecium* could result from selection acting on the joint functional activity of duplicate genes, such that eventual variation in expression and functional space might lead to the degeneration of one of the duplicates to a point where their contribution is negligible (Johri et al. 2022). Once this expression/functional contribution threshold is passed, the copy can be lost. One of the paths to expression divergence and consequently to gene loss could be the differential alteration of the core promoter region between paralogous genes.

Thus, the streamlined macronuclear genomes of *Paramecium* species that have undergone WGD and differential paralog loss present a system that facilitates comparative analysis of core promoter architecture and variation. We analyzed the transcription initiation regions (TIRs) across the genomes of two *aurelia* species (*P. tetraurelia* and *P. sexaurelia*) and their closest pre-WGD outgroup, *P. caudatum.* Three characteristics of the TIR (position, shape, and TIS site relative to a paralog) were identified and contrasted to expression level per gene per species, per two-paralog family for the *aurelias,* and for homologous genes between species (Figure 1). These comparisons serve as a means to explore the architecture of TIRs in streamlined genomes and assess whether core promoter erosion is a candidate mechanism underlying gene loss through divergence of transcriptional output between paralogs. Next, the regions surrounding focused TIR genes were surveyed for key eukaryotic core promoter motifs. Finally, the overarching architecture of *Paramecium* TIRs is compared to the TIRs of other eukaryotes.

**Figure 1.**
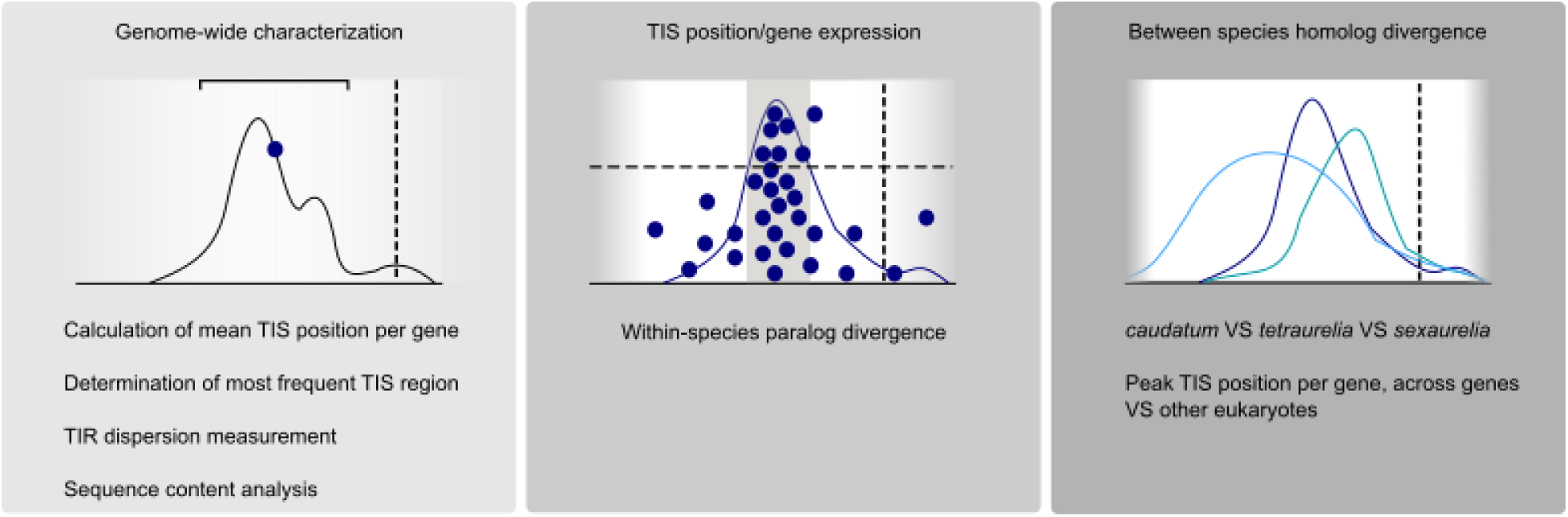
Cartoon representation of key concepts and analyses for this work. X axis represents the position in bp relative to the translation start site (vertical dashed line). Left, a transcription initiation region (TIR, black line) is determined per gene, and the mean transcription initiation site (mean TIS) is calculated (blue dot, see Methods). The dispersion of the TIR is also calculated (black bracket, see Methods). Middle, the mean TIS position per gene is compared to the gene’s expression. Position/expression profiles are binned by expression level (horizontal dashed line) and region (gray shading for region where most mean TISs occur). Position/expression profiles can be used to compare paralog genes in the *aurelias.* Right, the peak TIS per gene (see methods) is obtained per genome, and genome-wide TIS positions are then compared between *Paramecium* species and other eukaryotes.

## Results

### Transcription initiation occurs remarkably close to the translation start site in *Paramecium*, but differs between species

STRIPE-seq libraries for *P. caudatum, P. tetraurelia* and *P. sexaurelia* were trimmed for adapters and filtered to remove predicted rRNA sequences (see Methods). The filtered reads were then mapped to their corresponding genomes (Table S1). The regions between 150 bp upstream and downstream of the predicted translation start site per gene were extracted from the reference genomes (Aury et al. 2006, McGrath, 2014), and read starts were assigned positions per gene using custom scripts. Then, read start counts for uniquely mapped reads per position per gene per species were obtained, and their positions expressed relative to the translation start site. Only sites per gene per species that were present in replicate experiments (duplicate for *tetraurelia* and *sexaurelia,* triplicate for *caudatum*) and with a total coverage of ≥3 were kept for analysis. The area in bp covered by the total TISs per gene is considered the transcription initiation region (TIR) for that gene (Figure 1, left). Under this framework, the TIRs for >40% of the predicted coding genomes of the three species were obtained (Table 1).

**Table 1.**
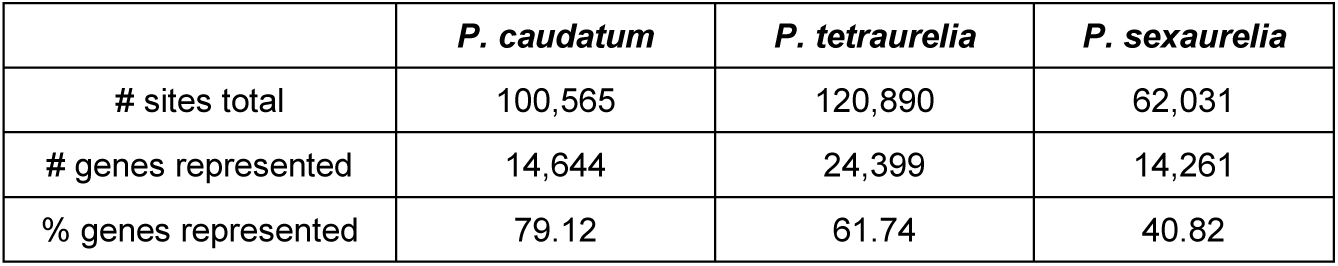
Number of TISs, genes, and percentage of genes (out of the total number of coding genes per species) for which there is at least one TIS identified.

In order to define the region where most TISs occur, the mean TIS position per gene was calculated (Figure 1, left). The distribution of mean TIS positions per species suggests that most genes initiate transcription within 50 bp of the translation start site (Figure 2A). A weighted prevalence score was applied to every site 150 bp up- and downstream of the translation start site (see Methods) in order to assess whether certain sites are preferentially used per species (Figure 2B). There is a large overlap of mean TIS position for the bulk of identified genes: the middle 50% region of all gene TISs in *P. caudatum* initiate transcription between -22 and -9 bp upstream of the translation start site, while for *P. tetraurelia* the TIS midspread lies between -26 and -11, and for *P. sexaurelia* between -25 and -11 (Figure 2). Despite the overlap in midspread TIS regions between species, the mean TIS by prevalence highlights the similarity in TIS usage between the *aurelias* compared to their outgroup (Figure 2B). Indeed, each species has significantly different distributions of mean TIS positions (pairwise Kolmogorov-Smirnov test with Bonferroni correction, p < 8.357×10^−9^), although these are relatively small. While statistically significant, *tetraurelia* and *sexaurelia* are minimally different from each other (KS statistic = 0.034), whereas *caudatum* is an outlier to both *aurelia*s (KS statistic = 0.166 and 0.185 vs *tetraurelia* and *sexaurelia*, respectively) (Figure 2B). The *aurelia* species tend to have longer intergenic spaces (median 161 bp in *tetraurelia,* 229 bp in *sexaurelia,* compared to 43 bp in *caudatum*) (Johri et al. 2017), which might account for the shift upstream of the translation start site in the *aurelias* relative to c*audatum.* While *caudatum* genes appear to have a marked preference for initiation at -9 with a secondary peak at -18, the aurelias have roughly the same prevalence peaks at -9, -11, and -18 bp (Figure 2B). These results suggest that TIRs have differentiated between species but still share a location constraint consistent with the reduced intergenic regions between genes and transcriptional streamlining.

**Figure 2.**
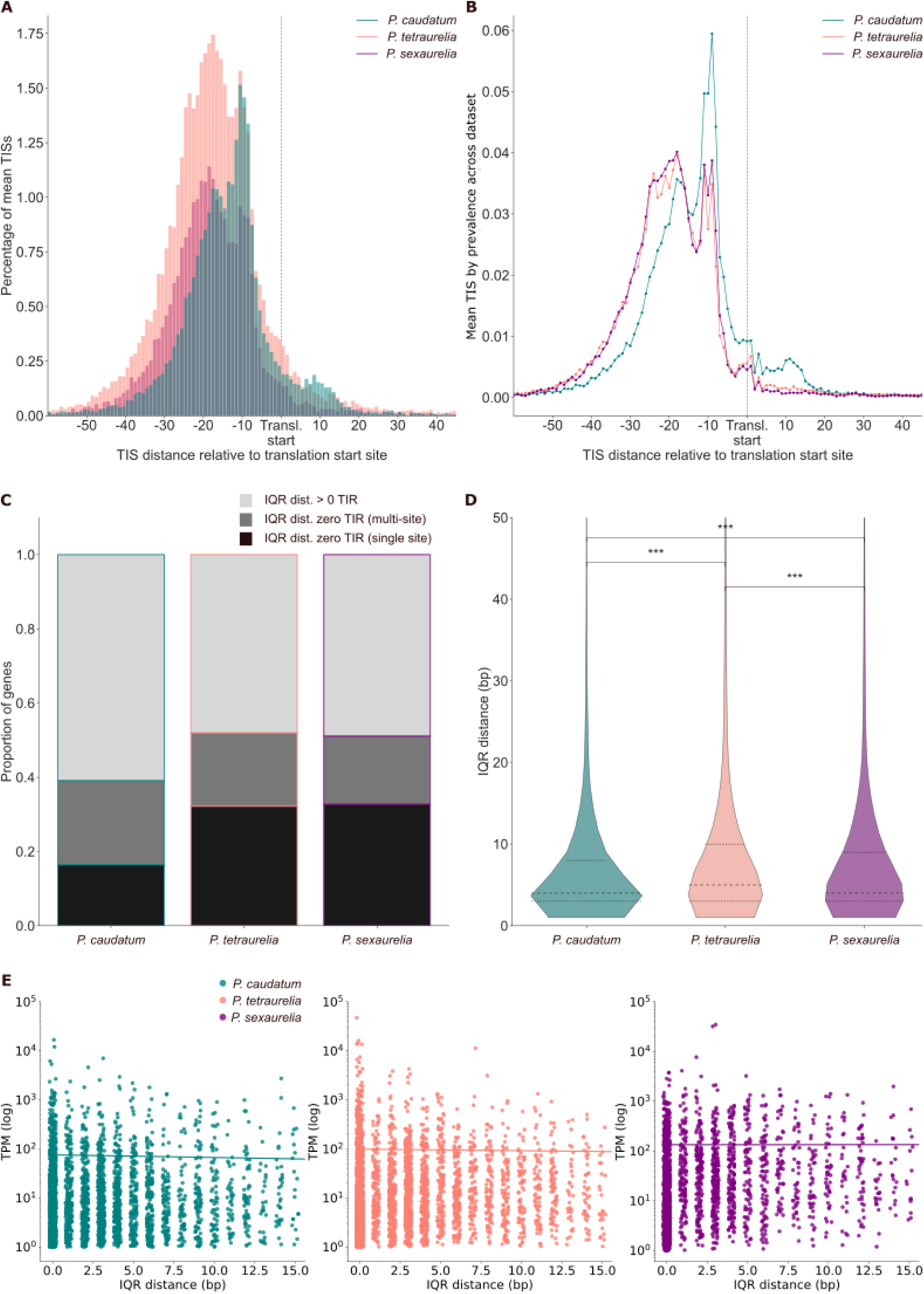
Genome-wide analyses of transcription initiation regions (TIRs) for three *Paramecium* species. **A.** Distribution of mean transcription initiation site (mean TIS) positions per species for the window of 60 to 50 bp up-and downstream of the translation start site. **B.** Mean TIS position by prevalence (see Methods) for the window of 60 to 50 bp up-and downstream of the translation start site. Note the ‘upstream shift’ that is common to the *aurelias*, but not to *caudatum.* **C.** Proportion of genes with different dispersion, measured by interquartile range distance in bp (see Methods), across their TIR shape per species. **D.** IQR distance of TIRs with dispersion > 0 (light gray in panel C) across the three species. The dotted lines represent the first and third quartile distances in bp, and the dashed lines represent median IQR distance. IQR distances for dispersed TIRs are significantly different between *P. caudatum, P. tetraurelia,* and *P. sexaurelia* (*caudatum* vs *tetraurelia—*left*, caudatum* vs *sexaurelia—*center, and *tetraurelia* vs *sexaurelia—* right, p < 6.013×10^−5^, see Methods). **E.** IQR distance versus gene expression and its linear regression model fit per species. Gene expression is independent of IQR distance.

### Transcription initiation regions are narrow across *Paramecium* and their dispersion is independent of gene expression intensity

Eukaryotes in general appear to have two general shapes to their TIRs, focused or dispersed. Thus, the dispersion of TIRs for *caudatum, tetraurelia,* and *sexaurelia* was measured using the interquartile range distance in bp per gene per species (IQR distance, see Methods). For all three species, the median TIR IQR distance is narrow: two bp for *caudatum* and zero bp for the *aurelias*. These results stem from the fact that there is a significant number of genes for which there is a single TIS comprising the TIR. For *caudatum,* 2,408 genes (16.44% of our dataset) are single-TIS, while for *tetraurelia* and *sexaurelia* 7,684 (31.49%) and 4,672 (32.76%) genes are single-TIS, respectively.

Moreover, among the genes with multiple TISs comprising their TIR, a significant fraction has the bulk of initiation (as measured by normalized read counts) occurring at a single site, such that their IQR distance remains zero. For *caudatum*, 3,318 genes (22.66% of the dataset) fall into this category, whereas 4,829 (19.79%) and 2,616 (18.34%) of the genes in *tetraurelia* and *sexaurelia* respectively have IQR distance of zero while having multiple TISs (Figure 1, left; Figure 2C). While the *aurelias* have comparable fractions of genes with IQR distance = 0 (51.28% for *tetraurelia* and 51.1% for *sexaurelia*), *caudatum* has a significantly lower proportion (39.1%, pairwise Mann-Whitney U test, p = 2.54×10^−74^ vs *tetraurelia* and p = 1.81×10^−62^ vs *sexaurelia*) (Figure 2C).

Among the genes with non-zero dispersion, the median IQR distance for *caudatum* and *sexaurelia* TIRs is 4 bp, and 5 bp for *tetraurelia* (Figure 2D, middle dashed lines). The middle 50% of the dispersed TIRs have an IQR distance of 8 bp for *caudatum,* 10 bp for *tetraurelia,* and 9 bp for *sexaurelia* (Figure 2D, lower and uppermost dashed lines). As a whole, these results show that the bulk of transcription-initiation events per gene across *Paramecium* occur within an area of 10 bp or less.

The dispersed TIR shape has been associated with highly expressed and/or constitutively expressed genes in a variety of organisms (Juven-Gershon and Kadonaga, 2010). Therefore, a linear regression was performed to evaluate whether IQR distance could predict gene expression (measured as log_10_ transcripts per million, see Methods) per *Paramecium* species. A total of 3,185, 4,138, and 2,431 genes with sufficient expression were used for *caudatum, tetraurelia,* and *sexaurelia*. Overall, the linear fit per species did not explain the variance in transcripts per million (TPM) (R^2^ < 0.013) as TIR dispersion increases (Figure 2E). These results suggest that TIR shape can change across genes without impacting gene expression across *Paramecium*.

### Highly expressed genes have a stronger constraint to be within a frequently used TIS region relative to the translation start site

Despite small (but significant) differences, most genes in *P. caudatum, tetraurelia* and *sexaurelia* have their mean TIS position within an area smaller than 50 bp (Figure 2A, B). Our data support previous work showing that the intergenic region in *caudatum* (median length 43 bp) is under strong constraint (Johri et al. 2017). While the constraint is less pronounced in the *aurelias*, which have longer intergenic regions (161 bp and 229 bp in *tetraurelia* and *sexaurelia* respectively), it is possible that despite the added genomic real estate TIRs are still limited to a somewhat well-defined region.

While TIR dispersion is unrelated to gene expression, the TIR position relative to the translation start site might bias the expression profile of its corresponding genes. With this in mind, we compared mean TIS position to expression per gene per *Paramecium* species. Gene expression in somewhat comparable levels (up to 10^2^ TPM) occurs for genes with mean TIS positions -150 to 150 bp from the translation start site (Figure 3) and is concentrated within the previously described most frequently used region (Figure 1, middle; Figure 2A, B). However, higher expression (>10^2^ TPM) genes appeared to be more concentrated within an area similar to the main TIS region (Figure 3A). In order to assess whether mean TIS position and high gene expression are linked, a Gaussian kernel density estimate (KDE) was used to approximate the distribution of the top 10% expressed genes over the full range of TIS positions relative to the translation start site (see Methods). Using a threshold of 40% of the KDE peak, a core region of 39-40 bp was defined, which included 91.8%, 91.1% and 87.7% of the mean TIS for high-expression genes in *caudatum, tetraurelia* and *sexaurelia* respectively (Figure 3A).

**Figure 3.**
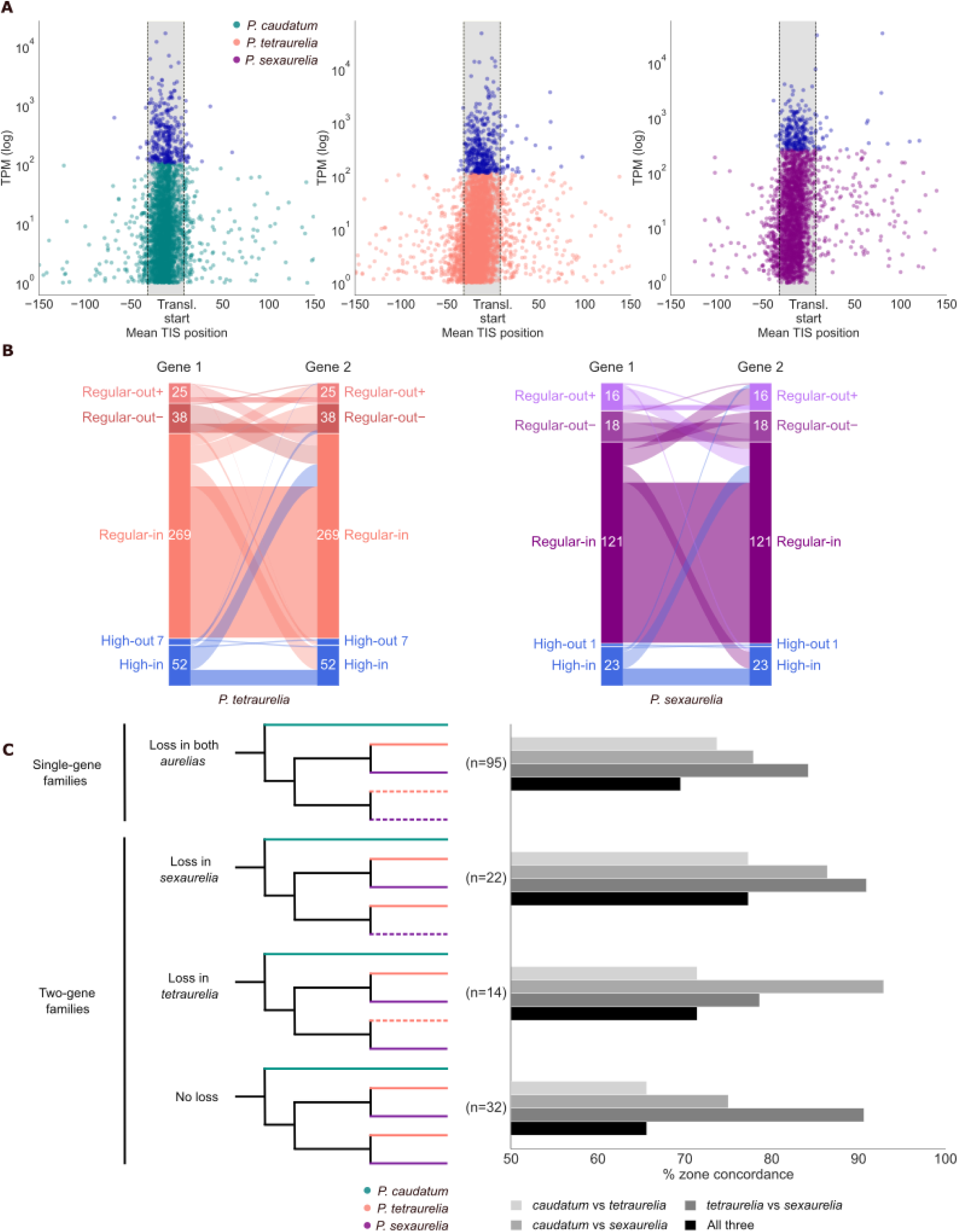
Gene expression is lightly impacted by conserved TIS position across gene families. A. Highly expressed genes (top 10% expression, blue) tend to be confined to a core region of about 40 bp upstream of the translation start site (gray shading) in *P. caudatum, P. tetraurelia* and *P. sexaurelia.* Note that high vs regular expression and in-core region vs up – or downstream of it (blue versus color and shaded area versus not shaded areas) generate the expression/position bins used for the datasets in panels B and C. B. Quantification of expression/position bins for two-gene families in *tetraurelia* (left) and *sexaurelia* (right). C. Inter-species comparison of expression/position profiles for homologs in single or two-gene families. Left section represents the number of members in a family, and dashed lines represent loss of a paralog. Right section is the corresponding percentage of gene families where the homologs share expression/position profiles in pairwise comparisons or for all three species.

The ∼40 bp area in which most highly expressed genes have their mean TIS includes most genes, regardless of expression, across the three species. Thus, Fisher’s exact tests were conducted to assess whether highly expressed genes are overrepresented in that region. Indeed, high-expression genes are between 1.41 and 2.25 times more likely to be within -32 to 8 in *tetraurelia* and *caudatum*, and -33 to 6 in *sexaurelia* (Figure 3A) (p = 0.055 per species, cumulative p = 1.92×10⁻⁷). The enrichment was strongest for *tetraurelia,* then *caudatum*, and weakest for *sexaurelia.* Next, Mann-Whitney U tests were done to test whether gene expression is in general higher for genes with mean TIS positions within the corresponding ∼40 bp region per species. For all species, even after excluding the top 10% expressed genes, genes with a mean TIR position within the ∼40 bp area have higher expression than genes outside of it (p=7.28×10⁻⁴, rank-biserial correlation r=0.107, p = 5.23×10⁻⁹, rank-biserial correlation r = 0.141, and p = 0.032, rank-biserial correlation r = 0.062 *caudatum, tetraurelia* and *sexaurelia*, respectively; combined p=5.71×10⁻¹¹). These results suggest that the mean position of a TIS relative to the translation start site has a small but significant role in increasing gene expression overall, but especially for highly expressed genes. This region is similar across species, despite the overall shift in TIR position when all genes are accounted for in the *aurelias* (Figure 2A, B).

### Among the *aurelias,* paralogs tend to have better conserved TIS mean position than expression level

From the above results we know that expression and mean TIS position per gene are linked, such that high-expression genes are more constrained to a narrow region in close proximity and upstream of the translation start site, and regular-expression genes might have their main TIS position vary. The spectrum of TIS position versus gene expression can be summarized, then, as follows: high-expression-in-region, high-expression-out-of-region, regular-expression-in region, and regular expression-out-of-region (Figure 1, middle). Given that gene expression is one of the predictors of gene loss in *Paramecium* (Johri et al. 2022), we reasoned that the distribution of gene family size in *tetraurelia* and *sexaurelia* might differ depending on the expression-TIS zones of its genes.

In *tetraurelia,* 4,138 gene families were tested for number of member genes versus their expression-TIS zones, of which retention occurred the most for highly expressed genes: 28.74% of highly expressed genes have at least one paralog, compared to 18.78% in regular-expression genes. In *sexaurelia,* a similar pattern exists, as 24.59% of highly-expressed genes have at least one paralog, whereas 15.27% of regular-expression genes retained any paralogs. These results paint a nuanced picture in which highly expressed genes are mostly constrained to initiating transcription within the previously described ∼40 bp region (Figure 3A, center and right), and the region itself is associated with higher gene expression overall, but it is expression intensity itself that changes the odds of paralog retention. This ultimately results in a non-random distribution of gene family sizes as a function of expression of at least one member paralog (chi-square test, p < 1.15×10^−4^), in agreement with previous work (Johri et al. 2022). While highly expressed genes retain more paralogs, this does not guarantee that the paralogs share expression profiles and/or mean TIS position. Hence, we compared the frequency of paralogs belonging to the same expression-TIS zone across gene families in *tetraurelia* and *sexaurelia.* Genes within the main TIS zone have higher retention of a paralog within the same zone (e.g. a regular-expressed in-region genes almost always have paralogs that are also regular-expression in-region) (Table 2). However, the paralogs may or may not have the same expression profile if one of the copies is highly expressed (Table 2).

**Table 2.**
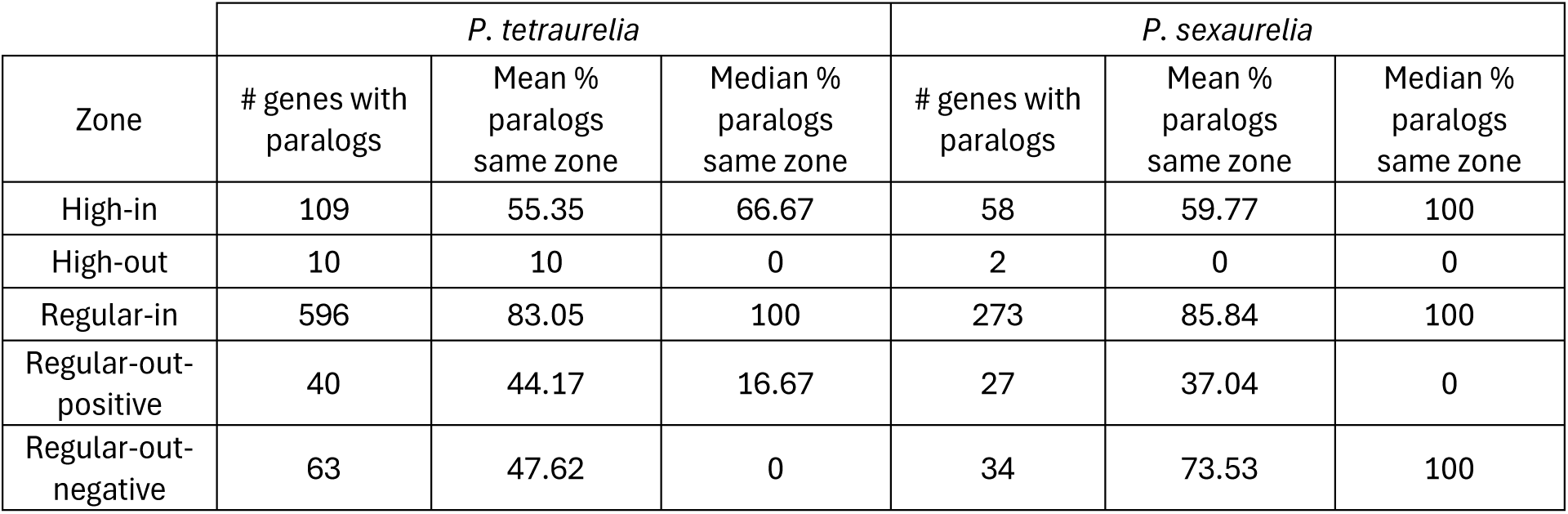
Paralog retention for expression level and mean TIS position region across two *Paramecium* species.

For both species, over 90% of multi-gene families for which we have complete TIS and expression data are comprised of 2 genes (310 and 149 families for *tetraurelia* and *sexaurelia* respectively). Thus, two-gene families were selected to assess in greater detail to what extent and in what way paralog TIS position and expression are diverging per species. Most two-gene families are zone-concordant (75.08% and 79.87% of families in *tetraurelia* and *sexaurelia,* respectively), driven by regular-expression-in-region genes, which comprise the most abundant category (Figure 3B), and then by high-expression genes. For the latter, 38-41% of high-expression-in-region genes have a sister gene with high expression and similar mean TIS position. Divergence in high-expression-in-region genes occurs mostly in expression level, as opposed to migrating mean TIS position out of the most frequently used (Figure 3B). Divergence in terms of TIS position but not expression is rare but differs between the species. For *tetraurelia*, more families have regular-expression-in-region genes with paralogs having mean TIS farther upstream from the translation start site, whereas in *sexaurelia* more regular-expression-in-region genes have paralogs with mean TIS closer, or even downstream, of the translation start site. Families with highly expressed genes outside of the region were too few to make meaningful comparisons, but it is worth noting that in the *aurelias* they appear to have a bias towards downstream TIS positions (Figure 3A). In light of these results, it is possible that upon undergoing WGD, most genes conserved the position of their ancestral TIR, and their expression then began to vary mainly as a function of modification of other core promoter elements. In both *aurelias*, over 93% of the two-gene families comprise sister paralogs, which makes this scenario likelier. So long as the core promoter retains its key elements, the impact of where the TIR is relative to the translation start site is small.

### TIR zone is conserved for most homologous single and two-gene families across *Paramecium*

The ancestor to the *aurelias* underwent two WGD events (McGrath et al. 2014, Long et al. 2023, Ni et al. 2025). As such, there are up to 4 genes in *tetraurelia* and in *sexaurelia* that are orthologous to every *caudatum* gene. Given that the number of four-gene families for which TIR data was recovered was small, we analyzed single and two-gene families per *aurelia* species to try to shed light on the divergence of homologous gene TIRs.

Pairwise TIR zone comparisons were made for 163 gene families ranging from single gene (where all but one paralog was lost in the *aurelias*) to incomplete two-gene (where one of the *aurelias* has both paralogs but the other does not) to complete two-gene (where both *aurelias* kept two paralogs) that have TIR zone data (figure 3C) (see Methods). For all categories, between 65.62% and 77.27% homologs in all three species conserved their TIR zone (Figure 3C). The highest rates of conservation of TIR zone occurred between the *aurelias* (78.57% to 90.90%) (Figure 3C). One possible explanation for this pattern is that the ancestor to the *aurelias*’ genes already had slightly different combinations of mean TIS/gene expression profiles relative to *caudatum*, which remained conserved once *tetraurelia* and *sexaurelia* diverged.

### The TIR is an AT-rich region surrounded by coding and non-coding sequences with higher GC content

The transcription initiation region is part of the core promoter sequence. In eukaryotes, a number of sequence motifs upstream of the TIR such as TATA boxes, Inr, and the DPE aid in positioning the transcription machinery such that transcription itself can begin (Yang et al., 2007, Juven-Gershon and Kadonaga, 2010, Roy and Singer, 2015). In *Paramecium,* the median intergenic space is fairly different across species, at 43 bp in *caudatum,* 161 bp in *tetraurelia,* and 229 bp in *sexaurelia* (Johri et al. 2017). It is thought that *caudatum* intergenic regions are under strong constraint (Johri et al. 2017), which is in agreement with our findings that the TIR region is well defined and conserved across *caudatum, tetraurelia,* and *sexaurelia*.

Focused TISs are usually used to find sequence motifs (Juven-Gershon and Kadonaga, 2010); thus the 100 bp upstream to 25 bp downstream of the peak TIS position (126 bp total) per gene for genes with an IQR distance of zero (5,325, 10,273, and 4,574 sequences for *caudatum, tetraurelia* and *sexaurelia*, respectively) were obtained and characterized (see Methods). For all species, the GC content of the sequence adjacent to the peak TIS position (the TIR, see Methods) is significantly lower than that of the 126 bp (100 bp upstream of the translation start site to 25 bp downstream of it) that ought to include the core promoter and the TIR and the average genomic GC (Table 3). The shift in nucleotide makeup of the TIR is likely to be related to its role relative to the rest of the core promoter, and to the coding sequence downstream of it. However, the GC content 100 bp upstream TIR to 25 bp downstream of it is lower overall for the *aurelias* as compared to *caudatum.* Regardless of species, the peak TIS shows no obvious preference for any specific nucleotide (beyond it being A/T) (Table 3). However, there appears to be a bias for what nucleotides surround the main TIS itself (Table 3). As a means to elucidate some of the additional signals that comprise the core promoter in *Paramecium,* the TIR sequences and approximate core promoter sequences per species were queried using STREME to search for candidate motifs (see Methods).

**Table 3.**
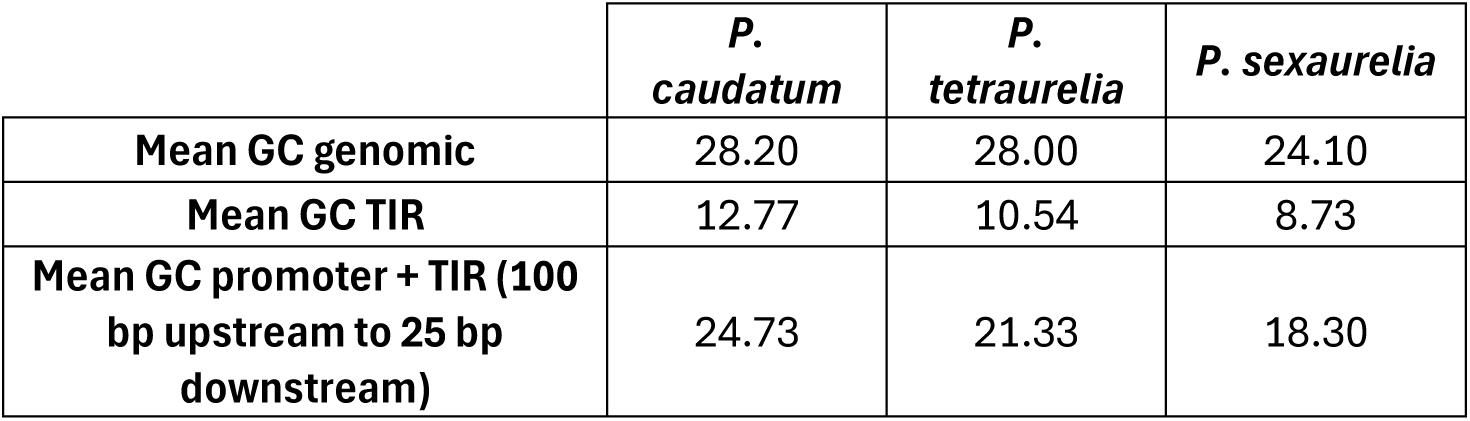
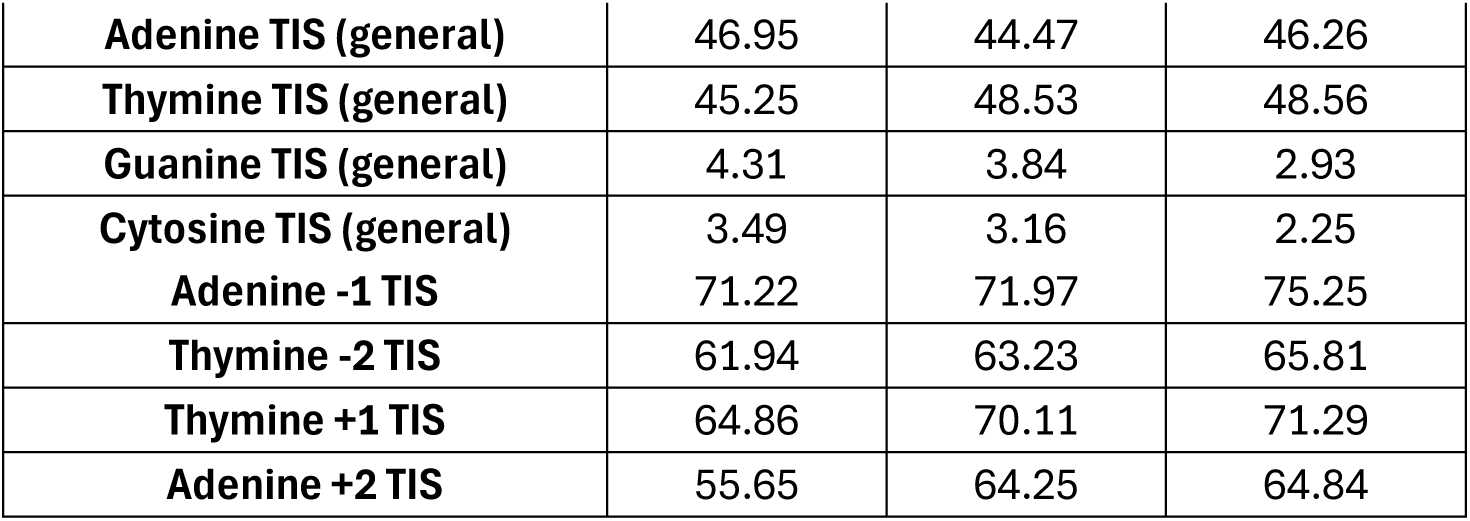
GC content and nucleotide preference expressed as percentages for TIR and adjacent region for three *Paramecium* species.

The Inr is among the conserved eukaryotic core promoter elements (Lo and Smale, 1996, Schumacher et al., 2003, Yang et al., 2007, Villao-Uzho et al., 2007, Roy and Singer, 2015), and in *T. vaginalis*, its consensus sequence is TCA_+1_T/CT/A (Schumacher et al., 2003). Thus, motif searches were performed for the TIR and the approximate promoter region (100 bp upstream to 25 bp downstream of the translation start site) for all genes with TIR data in *caudatum* and *sexaurelia.* While ATTCATT motifs were identified close to the TIS peak, in what would correspond to the Inr position, they were not well supported statistically. These results would suggest that the Inr is either strongly divergent from consensus or not a key element in the most focused promoters for *Paramecium.* Interestingly, the TCT motif (Parry et al., 2010), which is ribosomal and very similar to the Inr, was found for the three species. The *Paramecium* candidate TCT motif consensus was AAATCTTT, albeit also with low statistical support. Among the more upstream candidate motifs, repeated CTT motifs were found for all three species.

## Discussion

The *Paramecium* model system’s streamlined macronuclear genome, in conjunction with the differential resolution of paralog loss across the *aurelia* lineage, make for a valuable setup for the study of the evolution of transcriptional regulation. The expansion of the intergenic space in the *aurelias* is likely related to the small but conserved upstream shift that transcription initiation regions have across genes as a consequence of an increase in available sequence real estate relative to those of *caudatum* (Figure 2A, B). Expansion of TIRs, however, is modest, especially considering that *sexaurelia* TIRs have not shifted significantly despite median intergenic spaces being four times longer than the *caudatum* median intergenic region (Johri et al., 2017). While *caudatum* TIRs typically occur between -22 and -9 bp relative to the translation start site, -9 bp is clearly a site that is abundantly used in *caudatum*. For the *aurelias*, the shift in midspread region leaves -9 outside, which is matched by a more even usage of -9, -11, -18, and -24 in *tetraurelia* and *sexaurelia* (Figure 2D). The usage of -9 and -18 present the intriguing possibility that there might be a key distance between TIRs, perhaps determined by the molecular machinery in charge of transcription. Indeed, initial promoter melting requires a DNA unwound region of between 7 and 9 bp (Barnes et al. 2016). Moreover, 8 bp is thought to be the minimum necessary length for stabilizing transcription elongation in complexes consisting of RNA Pol II, oligonucleotides, and template DNA (Kireeva et al. 2000). In the *Paramecium* species used for this work, the median dispersed TIR width is at most 10 bp (in *tetraurelia*). Regardless of TIR shape, the area surrounding the mean TIS position is more AT rich than the surrounding core promoter and coding region. The lower GC content of a TIR could facilitate the crucial DNA unwinding and DNA-RNA hybrid length for reliable elongation during transcription. *P. caudatum* intergenic region sequences are severely constrained evolutionarily (Johri et al., 2017), but this does not appear to be as intense for the *aurelias*. The predicted intergenic lengths for other members of genus *Paramecium* place *caudatum* and *P. bursaria* as the species with the shortest average intergenic distances, as well as the smallest number of inferred ancestral WGD events (none for *bursaria*, one for *caudatum*) (Ni et al. 2025). Duplication of a TIR stretch or extension could then explain the shift to -18 bp for *tetraurelia* and *sexaurelia,* and perhaps for other species with a history of multiple WGD events; however, it is also possible that *caudatum* has undergone more extreme streamlining.

The TIRs across the three *Paramecium* species are in general terms narrow, as compared to those of other eukaryotes. Focused TIRs have been classified as having widths of up to 5 bp (Carninci, 2006, Miura et al., 2006, Main et al., 2013). These metrics would lead to an overall classification of TIRs in *Paramecium* as overwhelmingly focused. In this work, we used dispersion of the TIR as a metric for width without necessarily assigning focused or dispersed classifications. In *Paramecium,* TIR dispersion is not linked to differences in gene expression.

It is clear that for all three *Paramecium* species, there is a preferred translation-start proximal region within which the bulk of genes have their TISs (Figure 3A). Three insights are of special relevance here. First, this region has a weak but positive association to increased gene expression, even though it is expression itself that drives paralog retention in *tetraurelia* and *sexaurelia.* Second, neither mean TIS position nor shape are sufficient to explain expression divergence occurring to a greater or lesser extent across paralogs in *tetraurelia* and *sexaurelia.* Homologs between species often have similar mean TIS positions/expression profiles (Figure 3C). Since our inter-species analysis was limited to single- and two-gene families, a possibility is that the dynamics of TIR position and potential regulatory rewiring differ more in the larger but less numerous 3- or 4-gene families in *tetraurelia* and *sexaurelia.* While it is feasible that mean TIS shifts could happen with equal probability up- or downstream of the most frequently used TIS region, it is likely that a fraction of the downstream TISs are the result of misannotation, as downstream shifts are at risk of overlapping with coding sequence, which could result in synthesis of loss-of-function transcripts. Upstream shifts in the *aurelias* might also have occurred in part due to the increase in sequence real estate from expanded intergenic regions for many genes. Indeed, the ciliate model *Tetrahymena thermophila* has longer intergenic regions (5,500 bp on average), and its genes generate longer 5’ UTRs as well (192.54 bp) (Figure 4) (Ye et al., 2025).

**Figure 4.**
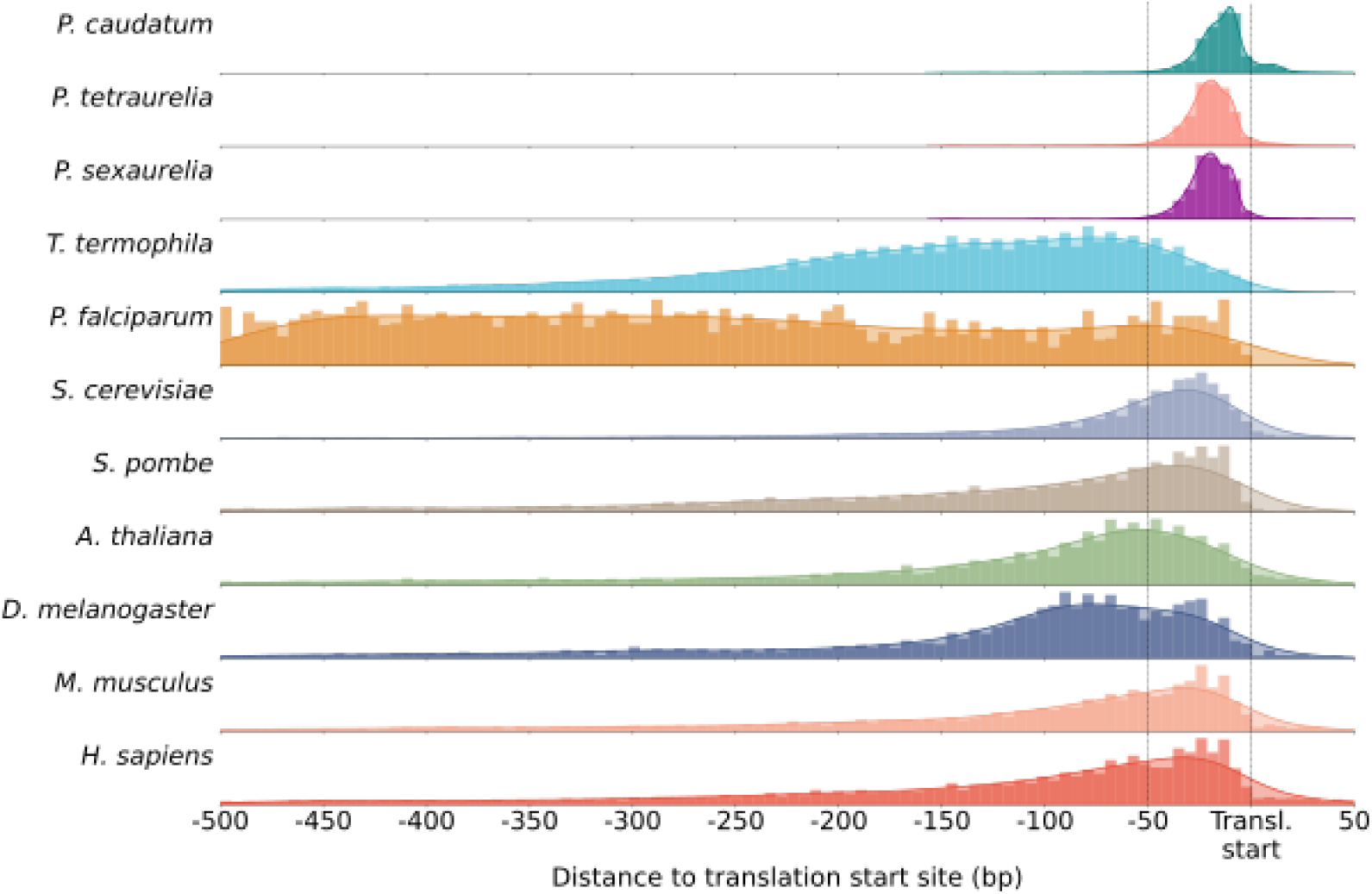
Density of TIS peak positions relative to the translation start site for eleven eukaryotic species. Note that for *Paramecium*, TIS peak positions are almost entirely within 50 bp upstream of the translation start site (between dashed lines).

While *Paramecium* has a predicted TATA-binding protein, which suggests that TATA boxes can be part of the core promoter of some of its genes, we found no obvious evidence of TATA-like sequences in the conventional -20 to -35 bp relative to the mean TISs across *caudatum, tetraurelia,* and *sexaurelia.* While a couple of motifs were identifiable within 50 bp upstream of the mean TIR across the three *Paramecium* species, nothing as obvious as the Inr in, for example, *T. vaginalis,* (Schumacher et al., 2003) was identified. Non-standard promoter architecture is also a characteristic of another unicellular eukaryote, *T. brucei* (Wedel et al., 2017), though this is in part because of the unique trypanosomal polycistronic transcription. Work in *Tetrahymena* suggests that it might have sequence signals that differ from those of metazoans for cleavage and polyadenylation (Ye et al., 2025). Moreover, it was proposed that *Paramecium caudatum* might have divergent Kozak sequences for translational initiation (Yamauchi et al., 1992). Thus, the identification of transcription factor binding motifs in the context of the compressed promoter of *Paramecium* promises to shed useful knowledge on how key regulatory information can be conveyed in minimal space. Despite extensive editing at the transition from micronuclear to macronuclear genome, *Paramecium* often maintains synteny between its macronuclear genomes (Long et al., 2023). Given the minimal intergenic space available between certain genes, it is possible that core promoter elements for one gene are shared between genes or overlap with the 3’ UTR of the upstream neighbor. The deleterious loss of regulatory information that loss of synteny could bring might be one of the reasons for maintenance of synteny in the macronuclear genome.

The third insight from this work comes from putting the *Paramecium* TIR in the context of that of other eukaryotes (Figure 1, right, Figure 4). In human, mouse, and *Plasmodium falciparum* (Carninci, et al., 2006, Adjalley et al., 2016) TIRs, pyrimidine-purine dinucleotides delineate transcription initiation sites. In the case of *Plasmodium*, which is also AT rich, a strong decline in GC content was also found around the most active TISs. These results suggest that a combination of signals, including py-pu dinucleotides likely form a conserved signal for transcription initiation across eukaryotes. However, the sequence makeup across lineages likely influences what those dinucleotides are, and additional signals might be needed when high AT content is present.

Beyond the dispersion or lack thereof for the TIR, the region where TIRs mostly occur in *Paramecium* is truly minimal, and perhaps close to the smallest a eukaryotic organism can allow, leaderless transcripts notwithstanding. Independently of the mechanism for transcription initiation (for example, scanning in *S. cerevisiae* or anchoring as in *S. pombe,* mouse and human) and of GC content in the promoter region, diverse organisms have at least some local maxima of TIRs within 50 bp upstream of the translation start site. In the extreme case depicted by *Plasmodium*, the mode for TISs occurs 50 bp upstream of the translation start site (Adjalley et al., 2016). It has been proposed that the physical expansion of 5’ UTRs can result from nonadaptive mechanisms (Lynch et al., 2005). Conversely, *Paramecium* species likely experience exceptionally efficient selection, which has likely driven the strong conservation in TIR position, dispersion, and GC content across species.

## Methods

### STRIPE-seq library preparation and sequencing

*Paramecium* cells were grown in up to 4 L Wheat Grass Powder medium (Pines International) prior to harvest. Cultures of *P. tetraurelia* strain 51 and *P. sexaurelia* strain AZ8-4 for harvest were separated from food bacteria by filtration on a 10 *µ*m Nitex membrane, and cultures of *P. caudatum* isolate 22-28 were separated by filtration on a 15 *µ*m Nitex membrane and. RNA was isolated using TRIzol (Ambion) and the manufacturer’s suggested protocol for tissue culture cells. STRIPE-seq libraries were generated as per Policastro et al., 2020.

Libaries were prepared in triplicate for *P. tetraurelia* and *P. sexaurelia,* and in quintuplicate for *P. caudatum.* These libraries were then sequenced with Illumina single-end 150 nt reads. STRIPE-seq reads were trimmed for adapter sequences and quality using Trimmomatic version 0.33 (Bolger, et al., 2014). Trimmed reads lacking the spacer (TATA) and 8-bp unique molecular identifier (UMI) sequences were filtered put using custom PERL scripts. Predicted ribosomal RNA sequences were obtained using RNAMMER version 1.2 (Lagesen et al., 2007). Read mapping was performed with Hisat2 version 2.1.0 (Kim et al., 2019). First, reads were mapped against the predicted RNAMMER rRNA sequences and filtered out. The remaining reads were then mapped to each corresponding genome: *P. tetraurelia* strain 51 V2 (Aury et al. 2006 et al.), *P. sexaurelia* strain AZ8-4 V2 (McGrath et al., 2014), and *P. caudatum* strain 34c3d (McGrath et al., 2014). Selection of candidate transcription initiation sites and regions was performed using SAMTools version 1.8 (Danecek et al., 2021), bcftools version 1.9 (Danecek et al., 2021), and custom scripts.

### Selection of candidate transcription initiation sites and regions (TIS and TIR)

Only uniquely mapped genes in a region from 150 bp upstream to 150 bp downstream of the predicted translation start site for coding genes across per replicate per species were used to search for candidate TISs. Read start counts per replicate were obtained and used to filter out unreliable candidate TISs. For the *aurelias*, which had data in triplicate, sites present in at least two of the three replicate experiments and with a sum coverage of 3 were kept as TISs per gene per species. For *P. caudatum*, sites present in at least three of the five replicates and with a sum coverage of 3 were kept as TISs per gene. Then, TISs read counts were averaged across replicates per gene per species and normalized to sum 1 to allow for intra- and inter-species comparisons.

A weighted mean prevalence score for the TISs per gene per species was used to determine the most frequent positions of TISs across *Paramecium*. For a given position, the grand mean of the normalized TIS read counts was calculated. Then, the number of genes with signal for that position was divided by the total number of genes with TIS data associated to them; this measured the prevalence of that TIS. The product of the grand mean signal and the prevalence was generated the mean prevalence score for every position (-150 to 150) per species. Pairwise differences between mean TIS site positions (*caudatum* vs *tetraurelia, caudatum* vs *sexaurelia, tetraurelia* vs *sexaurelia*) were tested using Kolmogorov-Smirnov testing and p-values were adjusted with Bonferroni correction.

Since a gene can have many TISs (and thus a TIR), mean TIS position per gene per species was calculated as follows: each TIS position was multiplied by its normalized count and then summed. The midspread TIS region per species was calculated as the area in bp between the position where 25% of mean TIS positions for all genes occurred, and the position where 75% of mean TIS positions occurred.

### TIR dispersion measurement

The interquartile range distance (IQR distance) in bp of TIRs per gene per species were obtained and used to measure TIR dispersion as follows: for every gene, its TISs were sorted by their distance to the translation start site. The cumulative sum of the normalized counts per TIS was calculated, such that the position relative to the translation start site at q1=25% of the counts and the position at q3=75% of the counts for that gene were obtained. The IQR distance then was position at q3 – position at q1. A three-way comparison of the transcriptome-wide IQR distance distributions for *caudatum, tetraruelia* and *sexaurelia* was performed via Kruskall-Wallis test. Given the high proportion of IQR distance = 0 genes, only the genes with IQR distance >0 were considered to have any dispersion, and were subjected to pairwise, two-sided Mann-Whitney U tests with Bonferroni correction to assess whether there are interspecies differences in how disperse TIRs are.

### Gene expression analysis

Given the fact that STRIPE-seq libraries are modified RNA-seq libraries, the length of trimmed and filtered reads was assessed to assess whether the reads could also be used as RNA-seq data or if they could also represent aborted transcription events. All reads had lengths greater than 35 bp and thus were considered appropriate to use as RNA-seq data for within-species gene expression analysis. The filtered and mapped reads used to identify TSSs were also processed as RNA-seq data. Transcript abundances in TPM were estimated using Stringtie 3.0.3 (Shinder, I. et al., 2026). The median TPM per gene across replicates per species was calculated. Only genes with median TPM >=1 were used for expression analysis and log_2_(median TPM+1) transformed in order to perform paralog expression comparisons using custom python scripts.

Linear regressions per species on IQR distance versus log_10_TPM across the transcriptome were performed, and Pearson’s r was used to evaluate the possibility of any correlation between dispersion and gene expression. Gaussian kernel density estimations (KDE) per species were performed using the top 10% expressed genes, a bandwidth of 10 bp (which is the most dispersed median distance in bp for TIRs across the three species), and a threshold of 40% of the KDE peak using custom python scripts. Next, the area (in bp) covered by the KDE was used to define a core region per species. One-sided Fisher’s exact tests were performed per species to test for overrepresentation of highly-expressed genes in the core region, and one-sided Mann-Whitney U tests were performed only on the bottom 90% expressed genes to test for whether genes within the core region have higher expression than genes outside of it using custom python scripts.

### Within-species paralog analysis and between-species homolog analysis

Gene families comprising *P. caudatum. P. tetraurelia* and *P. sexaurelia* were obtained from Gout et al. 2023. For the *aurelias*, gene families were queried for gene number per species relative to *P. caudatum* genes (up to 4 genes per species). The paralogs of genes with expression data were obtained and quantified using custom python scripts as a means to test for differential retention when at least one member of a gene family is highly expressed vs regularly expressed. Next, instances of genes with mean TIS + expression information where their paralog also had mean TIS + expression information were binned according to their expression and mean TIS position relative to the core expression region. This dataset comprised 818 families for *tetraurelia* and 394 families for *sexaurelia.* While 3 or 4 gene families with TIR data were recorded, they are so few that only 2-gene families were used for further analyses. For *tetraurelia,* 310 two-gene families were used; for *sexaurelia*, 149 two-gene families were queried. Comparisons between the two paralogs per family per species were done for their expression/position bin were performed and quantified using custom python scripts.

Two-gene families where both members had the same expression/position bin in *tetraurelia* and *sexaurelia* were used to search for their homologs in the remaining species. Families that had expression, TIR, and bin information in the three species, and that belonged to single or two-gene families in the *aurelias* were then subjected to pairwise comparisons (*caudatum* vs *tetraurelia, caudatum* vs *sexaurelia, tetraurelia* vs *sexaurelia*) to test whether homologs were concordant in their expression/position bin or not using custom python scripts.

### GC content calculation and motif search

Sequences for TIR and core promoter regions for genes with IQR distance = 0 for the three species were extracted as FASTA files with custom python scripts. Per species, the TIR sequence area was defined based on the median IQR distance in bp for IQR > 0 genes (8 bp up- and downstream of the peak TIS in *caudatum*, 10 bp for *tetraurelia,* and 9 bp for *sexaurelia*). The approximate core promoter was obtained by taking the area 100 bp upstream and 25 bp downstream of the peak TIS per gene. Ribosomal genes were obtained from the *P. tetraurelia* V2 annotation, and their homologs in *caudatum* and *sexaurelia* obtained from the gene family dataset from Gout et al., 2023. GC content calculation was performed using Biopython’s SeqUtils package. Motif searches were performed using the MEME suite v. 5.4.1 (Bailey et al., 2015) STREME tool, set to DNA and to search for a motif with minimum size 4 and search for 20 motifs.

### TIS comparisons to other eukaryotes

CAGE data for human and mouse was obtained from the FANTOM5 project (Kawaji et al., 2017). Start-seq data for *D. melanogaster* was obtained from GEO accession GSE96922 (Meers et al. 2017). CAGE data for *C. cerevisiae* was obtained from YeasTSS (McMillan et al., 2019) for the log phase in YPD medium condition (Lu and Lin, 2019). *S. pombe* CAGE data was obtained from YeasTSS (McMillan et al., 2019) grown in YPD medium (Thodberg et al., 2019). PEAT data for *A. thaliana* was obtained from Morton et al., 2014. Distances between the TISs and the translation start were calculated based off annotation files (*H. sapiens* GENCODE V19, *M. musculus* NCBI37, Ensembl release 67, *D. melanogaster* BDGP5 V67, *S. cerevisiae* R64-1-1 V109, *S. pombe* ASM294 V2.26, and *A. thaliana* TAIR10). If many TISs per gene were identified, the most used ‘peak’ TIS was obtained used as representative per gene using custom scripts. In the case of *T. thermophila* and *P. falciparum,* the annotated genomes resulting from the corresponding studies (which had experimental identification of TISs) were used, and the starting position of the 5’ UTR was used as a proxy for the peak TIS. The distance between peak TIS positions relative to the annotated earliest translation start site available for that gene was calculated using custom scripts, and the number of genes with a TIS peak per position used for plotting the histogram and generating the density plots for the different species.

## Acknowledgements

This research was supported by the National Science Foundation Division of Biological Infrastructure under Grant DBI-2119963 and MCB-1412738, by the National Science Foundation Division of Environmental Biology under Grant Num. DEB1927159. We wish to acknowledge R. Taylor Raborn, Weibo Zhang, and Zhiqiang Ye for their early contributions to developing this project.

