## Supplementary material for "Survey of transcription initiation in the streamlined genomes of *Paramecium*": Table S1

|  | # reads (raw) | # reads filtered | % reads mapping once | # genes represented (no singletons) |
| --- | --- | --- | --- | --- |
| *tetraurelia* 1 | 29556514 | 27009617 | 38.65 | 30060 |
| *tetraurelia* 2 | 21044473 | 20121709 | 30.06 | 21963 |
| *tetraurelia* 3 | 21966718 | 21403318 | 38.5 | 25920 |
| *sexaurelia* 1 | 8653016 | 8418571 | 15.36 | 12547 |
| *sexaurelia* 2 | 8608537 | 8371948 | 18.82 | 12960 |
| *sexaurelia* 3 | 8929634 | 8691072 | 18.74 | 13343 |
| *caudatum* 1 | 29126340 | 27212513 | 53.79 | 15500 |
| *caudatum* 2 | 29090511 | 27046158 | 25.32 | 9760 |
| *caudatum* 3 | 33659376 | 31698309 | 53.25 | 15641 |
| *caudatum* 4 | 39208735 | 35771609 | 54.78 | 15673 |
| *caudatum* 5 | 15497042 | 10908875 | 48.83 | 12456 |

**Supplementary Figures and Tables**

Table S1. STRIPE-seq of three *Paramecium* species. First column represents each species and its replicate number. Second column represents total number of reads obtained from sequencing runs. Filtered read counts are compiled after trimming for quality, adapters, and removing predicted rRNA sequences. If a gene only had sites with coverage of 1, it was considered a singleton and filtered out (see Methods). The last column accounts for genes that had enough non-singleton TIS data to be analyzed.

| ***P. caudatum* top 50 expressed** | **IQR distance** | **TIR shape** | **Mean position** | ***P. tetraurelia* top 50 expressed** | **IQR distance** | **TIR shape** | **Mean position** | ***P. sexaurelia* top 50 expressed** | **IQR distance** | **TIR shape** | **Mean position** |
| --- | --- | --- | --- | --- | --- | --- | --- | --- | --- | --- | --- |
| PCAU.43c3d.1.G00200135 | 144 | dispersed | 19.60 | PTET.51.1.G0020469 | 9 | dispersed | -133.98 | PSEX.AZ8_4.1.G0190105 | 0 | focused | -14.72 |
| PCAU.43c3d.1.G00200136 | 83 | dispersed | -70.14 | PTET.51.1.G0230218 | 33 | dispersed | -73.38 | PSEX.AZ8_4.1.G0350134 | 110 | dispersed | -11.26 |
| PCAU.43c3d.1.G00200132 | 58 | dispersed | 106.33 | PTET.51.1.G0090011 | 0 | focused | 1.19 | PSEX.AZ8_4.1.G1240017 | 59 | dispersed | 12.97 |
| PCAU.43c3d.1.G00200133 | 139 | dispersed | -17.04 | PTET.51.1.G0400116 | 4 | dispersed | -27.96 | PSEX.AZ8_4.1.G0070270 | 3 | dispersed | -9.15 |
| PCAU.43c3d.1.G00020355 | 0 | focused | -22.10 | PTET.51.1.G0370083 | 9 | dispersed | -4.50 | PSEX.AZ8_4.1.G0690173 | 2 | dispersed | 2.43 |
| PCAU.43c3d.1.G00200134 | 33 | dispersed | -101.35 | PTET.51.1.G0610114 | 0 | focused | -21.02 | PSEX.AZ8_4.1.G1700017 | 12 | dispersed | 2.82 |
| PCAU.43c3d.1.G00200131 | 37 | dispersed | -113.48 | PTET.51.1.G1840043 | 0 | focused | -3.22 | PSEX.AZ8_4.1.G0190108 | 2 | dispersed | -14.97 |
| PCAU.43c3d.1.G00020306 | 3 | dispersed | -8.66 | PTET.51.1.G0890107 | 4 | dispersed | -34.41 | PSEX.AZ8_4.1.G0070235 | 2 | dispersed | -19.83 |
| PCAU.43c3d.1.G00660090 | 0 | focused | -16.18 | PTET.51.1.G0670042 | 2 | dispersed | 2.35 | PSEX.AZ8_4.1.G1710023 | 2 | dispersed | 7.29 |
| PCAU.43c3d.1.G00110052 | 1 | dispersed | -9.98 | PTET.51.1.G1550027 | 33 | dispersed | -111.18 | PSEX.AZ8_4.1.G0340280 | 98 | dispersed | 20.90 |
| PCAU.43c3d.1.G00790027 | 3 | dispersed | -20.19 | PTET.51.1.G0080248 | 0 | focused | -31.51 | PSEX.AZ8_4.1.G0690115 | 0 | focused | -3.01 |
| PCAU.43c3d.1.G00840007 | 1 | dispersed | 3.35 | PTET.51.1.G0260234 | 3 | dispersed | -12.07 | PSEX.AZ8_4.1.G0450273 | 76 | dispersed | 8.87 |
| PCAU.43c3d.1.G00490172 | 3 | dispersed | 0.080 | PTET.51.1.G1440102 | 3 | dispersed | -23.82 | PSEX.AZ8_4.1.G0340086 | 0 | focused | -16.29 |
| PCAU.43c3d.1.G00240171 | 53 | dispersed | 4.49 | PTET.51.1.G0510203 | 7 | dispersed | -12.90 | PSEX.AZ8_4.1.G1600072 | 40 | dispersed | -1.37 |
| PCAU.43c3d.1.G00790028 | 0 | focused | -20.50 | PTET.51.1.G0560051 | 9 | dispersed | 12.04 | PSEX.AZ8_4.1.G1510071 | 34 | dispersed | 47.03 |
| PCAU.43c3d.1.G00270098 | 11 | dispersed | -22.67 | PTET.51.1.G0480006 | 39 | dispersed | 0.86 | PSEX.AZ8_4.1.G0970007 | 12 | dispersed | 67.70 |
| PCAU.43c3d.1.G00040075 | 0 | focused | -25.39 | PTET.51.1.G0500069 | 2 | dispersed | -6.11 | PSEX.AZ8_4.1.G0090371 | 0 | focused | -15.72 |
| PCAU.43c3d.1.G00650054 | 6 | dispersed | -10.46 | PTET.51.1.G1580036 | 0 | focused | -32.05 | PSEX.AZ8_4.1.G0890173 | 0 | focused | -18.95 |
| PCAU.43c3d.1.G00310178 | 0 | focused | -12.83 | PTET.51.1.G0020470 | 3 | dispersed | -6.00 | PSEX.AZ8_4.1.G0300056 | 81 | dispersed | 52.47 |
| PCAU.43c3d.1.G00070152 | 0 | focused | -18.41 | PTET.51.1.G0220279 | 0 | focused | -0.86 | PSEX.AZ8_4.1.G0190331 | 1 | dispersed | -4.95 |
| PCAU.43c3d.1.G00460139 | 1 | dispersed | -11.84 | PTET.51.1.G0200018 | 0 | focused | -23.25 | PSEX.AZ8_4.1.G0980045 | 18 | dispersed | 18.36 |
| PCAU.43c3d.1.G00130263 | 4 | dispersed | -17.94 | PTET.51.1.G0380085 | 24 | dispersed | -12.61 | PSEX.AZ8_4.1.G0010501 | 0 | focused | -17.49 |
| PCAU.43c3d.1.G00200074 | 3 | dispersed | -5.93 | PTET.51.1.G0130131 | 3 | dispersed | -17.02 | PSEX.AZ8_4.1.G0950156 | 12 | dispersed | 4.09 |
| PCAU.43c3d.1.G00920037 | 0 | focused | -16.62 | PTET.51.1.G0240214 | 8 | dispersed | -25.33 | PSEX.AZ8_4.1.G0230031 | 3 | dispersed | 8.89 |
| PCAU.43c3d.1.G00320069 | 2 | dispersed | 106.95 | PTET.51.1.G1570128 | 4 | dispersed | -13.93 | PSEX.AZ8_4.1.G0920007 | 110 | dispersed | 16.76 |
| PCAU.43c3d.1.G00080063 | 0 | focused | -8.17 | PTET.51.1.G0450093 | 0 | focused | -17.70 | PSEX.AZ8_4.1.G0460066 | 3 | dispersed | -28.87 |
| PCAU.43c3d.1.G00220083 | 4 | dispersed | -2.15 | PTET.51.1.G1450006 | 0 | focused | -14.50 | PSEX.AZ8_4.1.G0870107 | 1 | dispersed | -14.66 |
| PCAU.43c3d.1.G00100094 | 5 | dispersed | -21.45 | PTET.51.1.G0670111 | 0 | focused | -8.19 | PSEX.AZ8_4.1.G0250180 | 0 | focused | -5.91 |
| PCAU.43c3d.1.G00300023 | 3 | dispersed | -29.43 | PTET.51.1.G0200201 | 0 | focused | -7.50 | PSEX.AZ8_4.1.G0360176 | 0 | focused | -16.48 |
| PCAU.43c3d.1.G00510148 | 2 | dispersed | -5.31 | PTET.51.1.G0750134 | 0 | focused | -12.17 | PSEX.AZ8_4.1.G0830047 | 0 | focused | -26.34 |
| PCAU.43c3d.1.G00260229 | 6 | dispersed | -7.79 | PTET.51.1.G0070103 | 4 | dispersed | -17.20 | PSEX.AZ8_4.1.G0670129 | 0 | focused | -23 |
| PCAU.43c3d.1.G00500089 | 2 | dispersed | -5.83 | PTET.51.1.G1170146 | 0 | focused | -13.14 | PSEX.AZ8_4.1.G1290036 | 0 | focused | -24.77 |
| PCAU.43c3d.1.G00690037 | 0 | focused | -7.57 | PTET.51.1.G0460139 | 0 | focused | -17.75 | PSEX.AZ8_4.1.G0660040 | 4 | dispersed | -24.80 |
| PCAU.43c3d.1.G00240072 | 3 | dispersed | -11.07 | PTET.51.1.G0550140 | 0 | focused | -26.16 | PSEX.AZ8_4.1.G0190260 | 22 | dispersed | 5.13 |
| PCAU.43c3d.1.G00250109 | 1 | dispersed | -21.11 | PTET.51.1.G0070232 | 1 | dispersed | -22.01 | PSEX.AZ8_4.1.G0480180 | 0 | focused | -11.13 |
| PCAU.43c3d.1.G00010366 | 4 | dispersed | -23.83 | PTET.51.1.G1190130 | 0 | focused | -20.71 | PSEX.AZ8_4.1.G0550160 | 4 | dispersed | -2.98 |
| PCAU.43c3d.1.G00080072 | 26 | dispersed | -19.87 | PTET.51.1.G1050126 | 0 | focused | -6.46 | PSEX.AZ8_4.1.G0430268 | 0 | focused | 33 |
| PCAU.43c3d.1.G00370080 | 2 | dispersed | -2.82 | PTET.51.1.G0210028 | 46 | dispersed | 16.09 | PSEX.AZ8_4.1.G0440109 | 4 | dispersed | -0.85 |
| PCAU.43c3d.1.G00550055 | 6 | dispersed | -1.91 | PTET.51.1.G0650043 | 4 | dispersed | -32.29 | PSEX.AZ8_4.1.G0460240 | 0 | focused | -22.51 |
| PCAU.43c3d.1.G00930073 | 2 | dispersed | -32.84 | PTET.51.1.G0120096 | 0 | focused | -11.16 | PSEX.AZ8_4.1.G0010334 | 28 | dispersed | 15.17 |
| PCAU.43c3d.1.G00130201 | 0 | focused | -10.84 | PTET.51.1.G0510134 | 0 | focused | 2.012 | PSEX.AZ8_4.1.G0890095 | 43 | dispersed | 17.29 |
| PCAU.43c3d.1.G00370142 | 5 | dispersed | -14.36 | PTET.51.1.G1180100 | 0 | focused | -16.42 | PSEX.AZ8_4.1.G0020066 | 0 | focused | -11.62 |
| PCAU.43c3d.1.G00060164 | 1 | dispersed | -14.33 | PTET.51.1.G0160087 | 42 | dispersed | 79.95 | PSEX.AZ8_4.1.G0060400 | 2 | dispersed | -8.55 |
| PCAU.43c3d.1.G00320108 | 3 | dispersed | -10.18 | PTET.51.1.G1630088 | 0 | focused | -38.08 | PSEX.AZ8_4.1.G0330211 | 12 | dispersed | -6.55 |
| PCAU.43c3d.1.G00240230 | 0 | focused | -25.23 | PTET.51.1.G0100097 | 0 | focused | -6.56 | PSEX.AZ8_4.1.G1320113 | 4 | dispersed | -9.98 |
| PCAU.43c3d.1.G01930003 | 1 | dispersed | -2.69 | PTET.51.1.G0860127 | 7 | dispersed | -25.33 | PSEX.AZ8_4.1.G1030042 | 0 | focused | -5.98 |
| PCAU.43c3d.1.G00010342 | 3 | dispersed | -11.27 | PTET.51.1.G1160124 | 0 | focused | -59.62 | PSEX.AZ8_4.1.G0330185 | 0 | focused | -17.31 |
| PCAU.43c3d.1.G00250010 | 0 | focused | -9.75 | PTET.51.1.G0230216 | 0 | focused | -22.01 | PSEX.AZ8_4.1.G0450083 | 0 | focused | -27.07 |
| PCAU.43c3d.1.G00580103 | 0 | focused | -19.73 | PTET.51.1.G0030065 | 0 | focused | -15.80 | PSEX.AZ8_4.1.G1530079 | 101 | dispersed | 2.57 |
| PCAU.43c3d.1.G00840050 | 0 | focused | -9.75 | PTET.51.1.G0410229 | 1 | dispersed | -8.98 | PSEX.AZ8_4.1.G1750052 | 0 | focused | -18.64 |

Table S2. Highly expressed gene IQR distance, TIR shape, and mean TIS position relative to the TSS for *P. caudatum, P. tetraurelia,* and *P. sexaurelia*.
